# Automatic analysis of skull thickness, scalp-to-cortex distance and association with age and sex in cognitively normal elderly

**DOI:** 10.1101/2023.01.19.524484

**Authors:** Junhao Zhang, Valerie Treyer, Junfeng Sun, Chencheng Zhang, Anton Gietl, Christoph Hock, Daniel Razansky, Roger M. Nitsch, Ruiqing Ni, the Alzheimer’s Disease Neuroimaging Initiative

## Abstract

Personalized neurostimulation has been a potential treatment for many brain diseases, which requires insights into brain/skull geometry. Here, we developed an open source efficient pipeline BrainCalculator for automatically computing the skull thickness map, scalp-to-cortex distance (SCD), and brain volume based on T_1_-weighted magnetic resonance imaging (MRI) data. We examined the influence of age and sex cross-sectionally in 407 cognitively normal older adults (71.9±8.0 years, 60.2% female) from the ADNI. We demonstrated the compatibility of our pipeline with commonly used preprocessing packages and found that BrainSuite Skullfinder was better suited for such automatic analysis compared to FSL Brain Extraction Tool 2 and SPM12- based unified segmentation using ground truth. We found that the sphenoid bone and temporal bone were thinnest among the skull regions in both females and males. There was no increase in regional minimum skull thickness with age except in the female sphenoid bone. No sex difference in minimum skull thickness or SCD was observed. Positive correlations between age and SCD were observed, faster in females (0.307%/y) than males (0.216%/y) in temporal SCD. A negative correlation was observed between age and whole brain volume computed based on brain surface (females -1.031%/y, males -0.998%/y). In conclusion, we developed an automatic pipeline for MR-based skull thickness map, SCD, and brain volume analysis and demonstrated the sex-dependent association between minimum regional skull thickness, SCD and brain volume with age. This pipeline might be useful for personalized neurostimulation planning.

## 1. Introduction

Brain stimulation has been a potential nonpharmacological therapeutic intervention for a range of brain diseases. Different neuromodulation methods have been developed using transcranial brain stimulation (TBS), transcranial ultrasound stimulation [1], transcranial direct current stimulation, transcranial electrical stimulation [2], transcranial magnetic stimulation, brain radiotherapy, etc. Increasing numbers of neuroimaging techniques, e.g., diffuse optical tomography [3], functional near-infrared spectroscopy [4], transcranial ultrasound microscopy [5], and photoacoustic imaging [6-8], have been developed and applied to human brain imaging. The skull compartment exerts a strong influence on the imaging and stimulation results, suggesting the need to accurately examine its geometry and alterations in the population. Thinning and geometric changes in the skull may also occur due to aging and disease. An earlier study showed the association between tomographic characteristics of the temporal bone and transtemporal window quality on transcranial Doppler ultrasound in patients with stroke or transient ischemic attack [9]. Skull shape abnormalities in ischemic cerebrovascular and mental diseases in adults have also been reported [10]. Optimization of the probe location on the head for neuroimaging/brain simulation thus has added value in various neurostimulation and imaging studies, such as simulation ultrasonic wave propagation and acoustic transmission [11-16].

Computed tomography (CT) imaging has been used to investigate the changes in bone thickness in 123 people (female/male) aged between 20-100 years [17, 18]. However, radiation exposure, particularly to the brain, is not ideal for volunteers or patients. Analysis using routine structural magnetic resonance imaging (MRI) data is a feasible alternative. Structural MRI-based automatic skull reconstruction from images was challenging, as compact bone has a very low signal in MRI. Recent efforts have enabled the development of accurate and automatic head segmentation/skull stripping pipelines from MRI for individualized head modelling [19, 20], which is not immediately applicable for skull analysis.

Here, we design a pipeline for automatically computing the skull thickness map, scalp-to-cortical distance (SCD), and brain volume based on structural T_1_-weighted (T_1_w) MRI data from the Alzheimer’s Disease Neuroimaging Initiative (ADNI 1 and 2) [21]. Cortical thinning has been found to be associated with amyloid-beta (Aβ and tau accumulation in Alzheimer’s disease (AD) and prodromal AD [22], as well as in clinically normal older adults [23], and associated with increasing age [24]. The changes in skull thickness and SCD with age and sex have not been evaluated in a large cohort, which is relevant in brain stimulation applications. To our knowledge, this study is the first to assess MRI-based skull thickness in a group of 407 cognitively normal older adults (71.9±8.0 years, 60.2% female).

## 2. Methods

### 2.1 MRI data

Data used in the preparation of this article were obtained from the ADNI database (adni.loni.usc.edu) [1]. The ADNI was launched in 2003 as a public□private partnership led by Principal Investigator Michael W. Weiner, MD. The primary goal of the ADNI has been to test whether serial MRI, positron emission tomography (PET), other biological markers, and clinical and neuropsychological assessments can be combined to measure the progression of mild cognitive impairment (MCI) and early AD. For up-to-date information, see www.adni-info.org. The dataset analysed includes 407 T_1_w MRI scans at 3 T of cognitively normal older adults between the ages of 51-100 years when scanned. Demographic information on the sex and age of cognitively normal older adults is presented in **Table 1**. The MRI imaging protocols of the ADNI study that were used to acquire the T_1_w MRI scans were described in [25] (field of view = 255×239×207 *mm*^3^, voxel size = 1 *mm*^3^). The majority of cognitively normal participants in the ADNI are of white ethnicity, with a mean education of 16 years and Mini-Mental State Examination score of 29.

**Table 1.** Comparison of different preprocessing methods for T1w MR segmentation with CT as the ground truth.

| **Pipeline** | **Skull** | **Scalp mask** | **Outputs** | **Subcortical analysis** | **Operation system** |
| --- | --- | --- | --- | --- | --- |
| BS | 0.7654 | 0.9779 | Label, mask, surface mesh | Yes | Windows/Unix |
| FSL | 0.7348 | 0.9639 | Label, mask | Yes | Unix |
| SPM | 0.7913 | 0.9743 | Probability map | No | Windows/Unix |

BS: BrainSuite Skullfinder preprocessing [26, 27], FSL: FSL Brain Extraction Tool [30, 31],
SPM: SPM 12-based unified segmentation [32].

### 2.2 MRI data analysis pipeline

The fully automated pipeline BrainCalculator was designed for processing T_1_w structural MR of the human brain (**Fig. 1a**). The BrainSuite (BS) 21a [26] skullfinder toolbox was used to segment the cortex, scalp, inner and outer skull surfaces based on a sequence of morphological transformations [27]. The initial skull stripping by using the skullfinder toolbox automatically computed the intensity threshold of the skull and scalp, which was then used to extract the skull and scalp surfaces with mathematical morphology. If the computed intensity threshold yields poor segmentation results, manual change of parameters is necessary for reaching higher accuracy. Next, the images were processed by a series of correction and masking algorithms to extract the cortex surface of the brain. Brain volume was calculated based on the cortex surface of the brain. The whole pipeline takes approximately 8 minutes for one dataset, as tested on Dell XPS 15 with Intel i7-9750H CP. After extracting the four surface meshes for the cortex, scalp, inner and outer skull surfaces from the MRI, 5×10^5^ points were uniformly sampled on the skull and scalp surfaces to form point clouds. The cortex surface was sampled with 5×10^6^ points, as the surface area is much larger than the skull and scalp surfaces. The SCD was calculated based on the cortex and scalp surface. The estimated skull thickness (is defined by the Euclidean distance between the two nearest points on the inner and outer skull surfaces). To eliminate outliers in point sampling, for each point on a surface, we searched for the 30 nearest points on the other surface and computed the root mean square as the thickness: 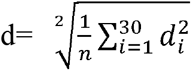 (where is *d*_*i*_ the Euclidean distance between the point of analysis and the *i*^*th*^ nearest point on the other surface)

**Fig. 1.**
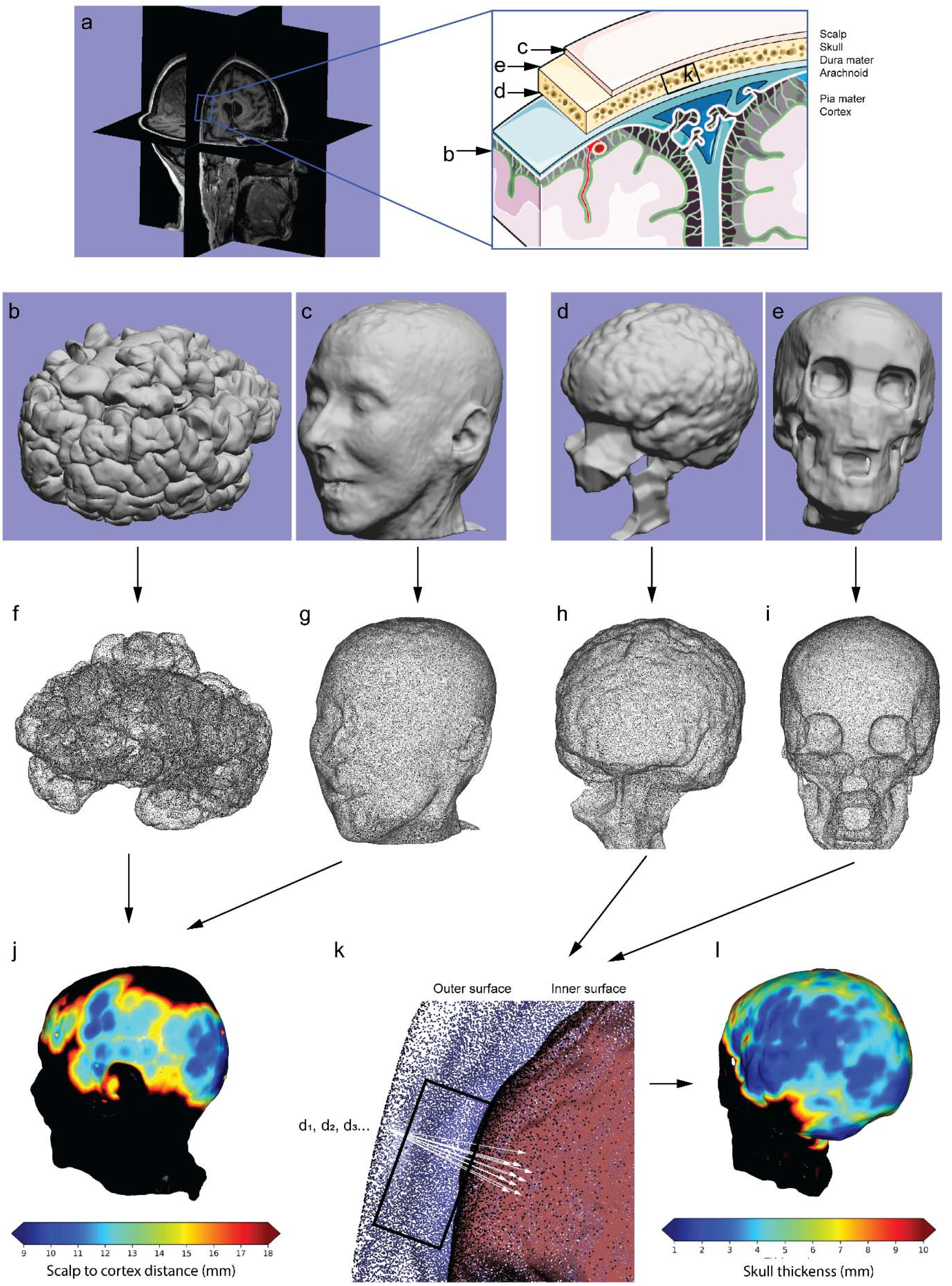
Analysis pipeline for human skull thickness and scalp-to-cortex distance (SCD) using T_1_w MRI data. (**a**) Original MRI file and zoom-in-view summarizing position of b-k; (**b-i**) Surfaces and corresponding point clouds for (**b, f**) cortex, (**c, g**) scalp, (**d, h**) inner skull and (**e, i**) outer skull; (**j**) SCD map. Scale bar = 9-18 mm (blue□red); (**k**) Zoomed-in view of the skull (in a) indicating the inner surface (red dots) and outer surface (blue dots). *di* is the *i*^*th*^ nearest point on the inner surface to each point on the outer surface (arrow line). (**l**) Skull thickness map. Scale bar = 1-10 mm (blue□red). Representative images based on T_1_w MR from one 79-year-old male.

To efficiently search for the nearest points, we used the K-dimensional tree algorithm to speed up the computation process [28]. The computation took approximately 3 minutes for the skull thickness map (**Fig. 1j**) and 5 minutes for the SCD (**Fig. 1l**). As the sphenoid, temporal, parietal, and occipital bones are important in brain stimulation studies, we chose calculations for these regions as examples. The points with minimum values in the regions of the sphenoid, temporal, parietal, and occipital bones can be directly located in the interactive visualization of the thickness map ((**Fig. 1j**). The user can choose a location on the skull with the cursor to retrieve the thickness value. Our opensource pipeline BrainCalculator is available at https://github.com/Junha0Zhang/BrainCalculator. The Open3D library in Python was used in our data processing [29].

### 2.3 Comparison of different preprocessing packages

With the aim of automatic computing, we compared the suitability and performance of different segmentation approaches without manual correction. We computed the skull meshes and thickness maps based on segmentation using BS Skullfinder [26, 27] and FSL Brain Extraction Tool (BET) 2 [30, 31] and Statistical Parametric Mapping (SPM) 12-based unified segmentation [32]. We compared the different approaches using a registered pair of T_1_w MR-CT images from the “Retrospective Image Registration Evaluation” dataset (Vanderbilt University) [33]. Detailed information on MRI and CT is described in [33]. We used 3D Slicer [34] to label the skull by thresholding from the CT scan as the reference. We then used flood filling to fill the unselected voxels inside the skull. For BET and BS, we segmented the MRI skull by subtracting the inner skull mask from the outer one. We computed the skull label directly by SPM12. We did not include the lower parts of the skull in the comparison, as they were either noisy or absent in the processing by all three methods. In addition to the skull, we computed the scalp mask (whole head) by thresholding. To compare the similarity of different analysis methods, we computed the Dice coefficients for each of the labels and the CT pair. The Dice coefficient measures the similarity of two sets of data and is defined as 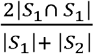, where |*S*| is the total number of voxels inside a skull label or scalp mask in our study.

### 2.4 Statistics

Normal distribution was determined by using the D’Agostino & Pearson test. For comparison of age between females and males, the nonparametric Mann□Whitney test was used. Two-way ANOVA with Bonferroni *post hoc* correction was used to compare the differences between groups by using GraphPad Prism (GraphPad, US, v9.0). Nonparametric Pearson correlation analysis was used to assess the association between age and SCD, skull thickness and brain volume for normally distributed data. All data are the mean ± standard deviation. Significance was set at p < 0.05.

## 3. Results

### 3.1 Development of the open-source pipeline BrainCalculator

We developed an open-source fully automated pipeline BrainCalculator for the computation of human skull thickness, SCD, and brain volume (**Fig. 1**). The original T_1_w MRI file of the human head (.nii) from ADNI (**Fig. 1a**) was processed by using BS Skullfinder to generate 3D meshes (.obj) for the cortical surface, scalp surface, and inner and outer skull surface (**Figs. 1b-e**). All datasets were registered to the same coordinate. Brain volume was calculated based on the cortex surface of the brain (**Fig. 1b**). Next, uniformly sampled points from the four surfaces for the cortex, scalp, inner and outer table (**Figs. 1b-e**) were used to generate corresponding point clouds (.pcd) (**Figs. 1f-i**). Earlier studies have used an exhaustive neighbor search of points [18]. To efficiently search for the nearest points, we used the K-dimensional tree algorithm to speed up the computation process [28]. For every point in the scalp and outer skull surface (**Figs. 1g, i**), the closest point in its paired data (cortex/inner skull) was identified (**Fig. 1k**). Next, we computed the skull thickness (**Fig. 1j**) and SCD maps (**Fig. 1l**) for all the selected points and regions. The skull thickness map is visualized, and the value is reported when the user selects the location on the skull by a cursor. The whole pipeline takes approximately 8 minutes for one dataset as tested on Dell XPS 15 with Intel i7-9750H CP.

### 3.2 Comparison of different preprocessing approaches

Next, we compared different open-source packages in the suitability and performance for automatic (without manual correction) skull segmentation and SCD computation. We demonstrated the utility of our pipeline based on BS Skullfinder preprocessing [26, 27] along with FSL Brain Extraction Tool (FSL) 2 [30, 31] and SPM12-based unified segmentation [32] for computing the skull meshes and thickness maps. SPM also provided probability maps for cerebral spinal fluid, white matter and gray matter, however did not provide scalp map. To evaluate the performance and compare the similarity of different analysis methods, we computed the Dice coefficients for each of the labels and the CT pair. CT data were used as ground truth. The Dice coefficient measures the similarity of two sets of data. Higher similarity of the segmentation to the CT reference is indicated by the closeness to 1.0. We did not include the lower parts of the skull in the comparison, as they were either noisy or absent in the processing. We found that the three methods BS, FSL, and SPM generated globally similar segmentations for the skull and scalp mask using the T_1w_ MRI data (**Table 1, Fig. 2**). However, mismatches in the temporal bone, sphenoid bone and occipital bone were observed in the automatically processed data based on FSL BET segmentation when overlaying the segmentation with CT (**Figs. 2c, g**). This mismatch could be observed in the skull thickness map generated based on FSL BET segmentation (**Figs. 2i, j**). Given the importance of the temporal bone, sphenoid bone and occipital bone regions in focused ultrasound stimulation and other neurostimulation approaches, we decided to use BS-based segmentation in our automatic pipeline.

**Fig. 2.**
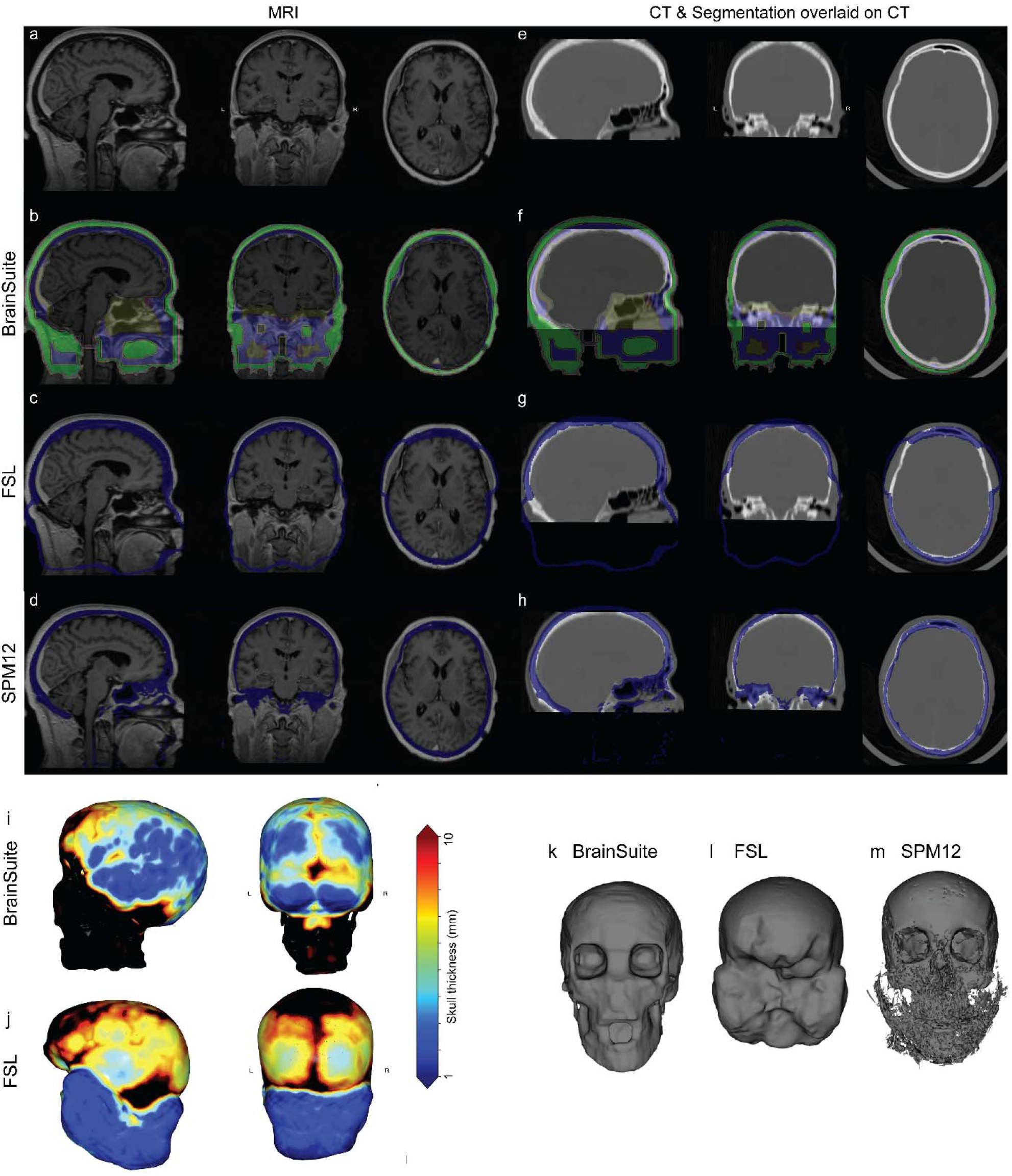
Comparison of different pipelines for skull and scalp-to-cortex map based on T_1_w MR-CT registered data. Images are displayed in sagittal, coronal and horizontal views. (**a, e**) T_1_w MR and CT image of human head; (**b, f**) BrainSuite skullfinder segmentation on MR and overlay of segmentation on CT: area between scalp and outer skull (green); area between inner and outer skull (blue); (**c, g**) FSL BET and (**d, h**) SPM12 segmentation on MR and overlay of segmentation on CT: area between inner and outer skull (blue); (**i, j**) Comparison of skull thickness map from BrainSuite and FSL. Scale bar = 1-10 mm (blue□red); (**k-m**) Comparison of skull mesh from FSL, BrainSuite, SPM12 processed using Braincalculator; Original MR and CT data source: “Retrospective Image Registration Evaluation” study [33].

### 3.3 Demographics

We collected data from the ADNI1 and ADNI2 databases from 407 cognitively normal older adults (71.93±7.98 years, 60.2% female). Detailed demographic information is presented in **Table 2**. Normal distribution analysis was performed by using the D’Agostino & Pearson test. The dataset assessed the normality test, with p values of 0.0760 (male, n = 162) and 0.3173 (female, n = 245), and the corresponding K2 values were 5.154 (male) and 2.296 (female). The ethnic background of the dataset from AD was white in majority.

**Table 2.**
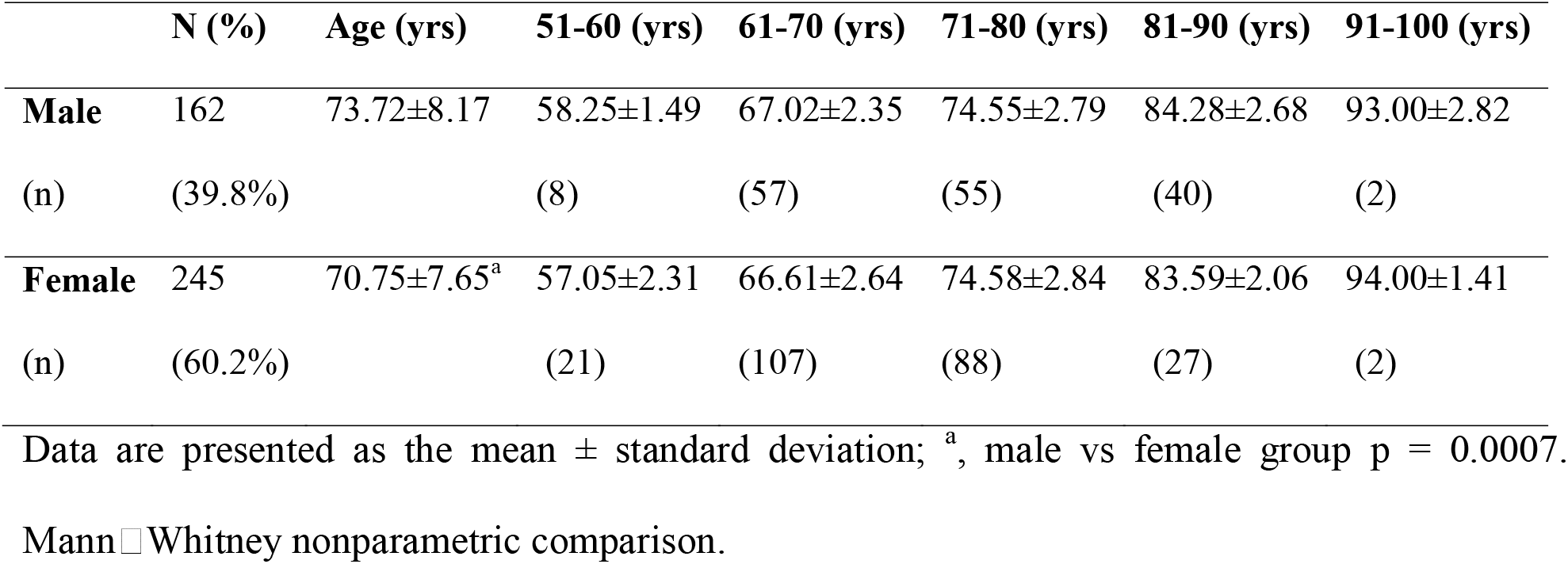
Demographic data of cognitively normal older adults in the study.

|  | **N (%)** | **Age (yrs)** | **51-60 (yrs)** | **61-70 (yrs)** | **71-80 (yrs)** | **81-90 (yrs)** | **91-100 (yrs)** |
| --- | --- | --- | --- | --- | --- | --- | --- |
| **Male** | 162 | 73.72±8.17 | 58.25±1.49 | 67.02±2.35 | 74.55±2.79 | 84.28±2.68 | 93.00±2.82 |
| (n) | (39.8%) |  | (8) | (57) | (55) | (40) | (2) |
| **Female** | 245 | 70.75±7.65<sup>a</sup> | 57.05±2.31 | 66.61±2.64 | 74.58±2.84 | 83.59±2.06 | 94.00±1.41 |
| (n) | (60.2%) |  | (21) | (107) | (88) | (27) | (2) |

Data are presented as the mean ± standard deviation; <sup>a</sup>, male vs female group p = 0.0007.
Mann-Whitney nonparametric comparison.

### 3.4 Association with age changes in skull thickness in the left hemisphere of male and female cognitively normal older adults

Next, we applied our analysis pipeline to compute the skull thickness map for the 407 T_1w_ MR datasets from ADNI (**Figs. 3a, b**). We chose the sphenoid, temporal, parietal, and occipital bones as examples to present the computed results for minimum skull thickness, as these brain regions are important in brain stimulation studies (**Table 2**). Group comparisons of the minimum skull thickness were performed between sex and age groups. There was a trend of higher skull thickness in male participants than in female participants, although the difference was not statistically significant. Significant skull thickening was found in the temporal bone of male participants in the 71- to 80-year-old group compared to the 60- to 70-year-old group (1.69±0.21 mm, n = 55 vs 1.59±0.19 mm, n = 57; p = 0.0402). In addition, significant skull thickening was found in the sphenoid bone of female participants in the 71- to 80-year-old group compared to the 51- to 60-year-old group (1.75±0.27 mm, n = 88 vs 1.53±0.23 mm, n = 21; p = 0.0037). In the sphenoid bone of female participants, there was a slight increase associated with age (r = 0.1734, p = 0.0066), with +0.202%/y. In other regions of skull bones, no correlation between age and skull thickness was detected in either sex. The magnitude of skull thickening was higher in the sphenoid bone and occipital bone of females than in those of males. In the temporal bone and parietal bone, the rates of change were similar in the female and male groups.

**Fig. 3.**
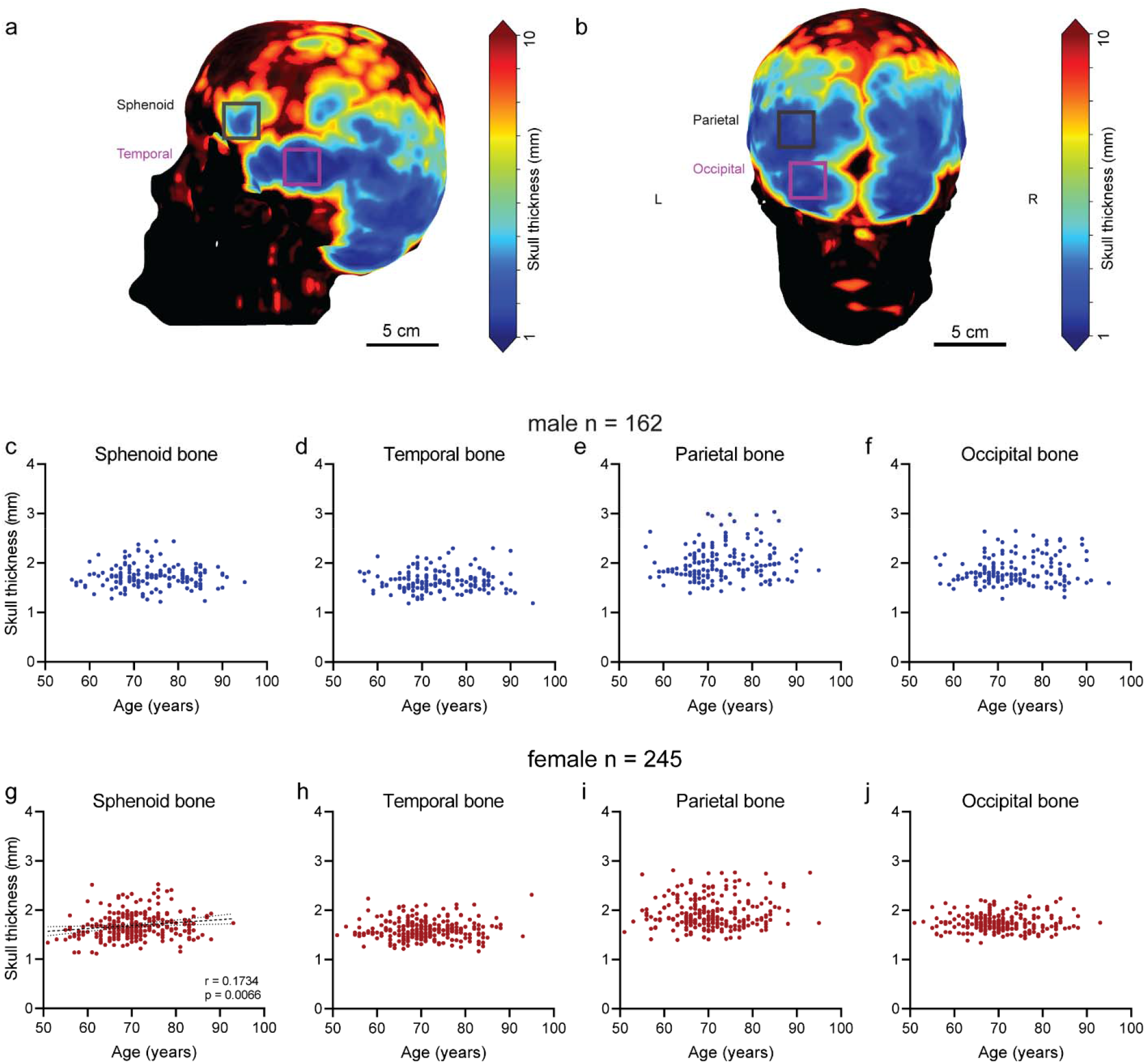
Skull thickness map and association between age and skull thickness in male and female older adults. (**a, b**) Representative skull thickness map-based T_1_w MR from one 55- year-old male; grey square indicates sphenoid bone (a) and parietal bone (b); magenta square indicates temporal bone (a) and occipital bone (b). Scale bar = 5 cm; Color scale = 1-10 mm (blue□red); (**c-f**) Association between age and skull thickness in male participants (n = 162); (**g- j**) Association between age and skull thickness in female participants (n = 245).

### 3.5 Sex-dependent age-associated increase in SCD in cognitively normal older adults

Next, we applied our analysis pipeline to compute the SCD map (**Figs. 4a, b**). Group comparison was performed between male and female groups of all ages and of different age groups and between different age groups of each sex (**Table 3**). No difference between male and female participants was observed. Significantly increased SCD was found in all four regions of both male and female participants in the older groups compared to the younger groups. We observed that in all the regions, there was an increased SCD associated with age in both the male and female groups. The slope of the increase in SCD was steeper in the female group (0.307%/y) than in males (0.216%/y) in the temporal cortex and was comparable in other brain regions (**Table 3**).

**Fig. 4.**
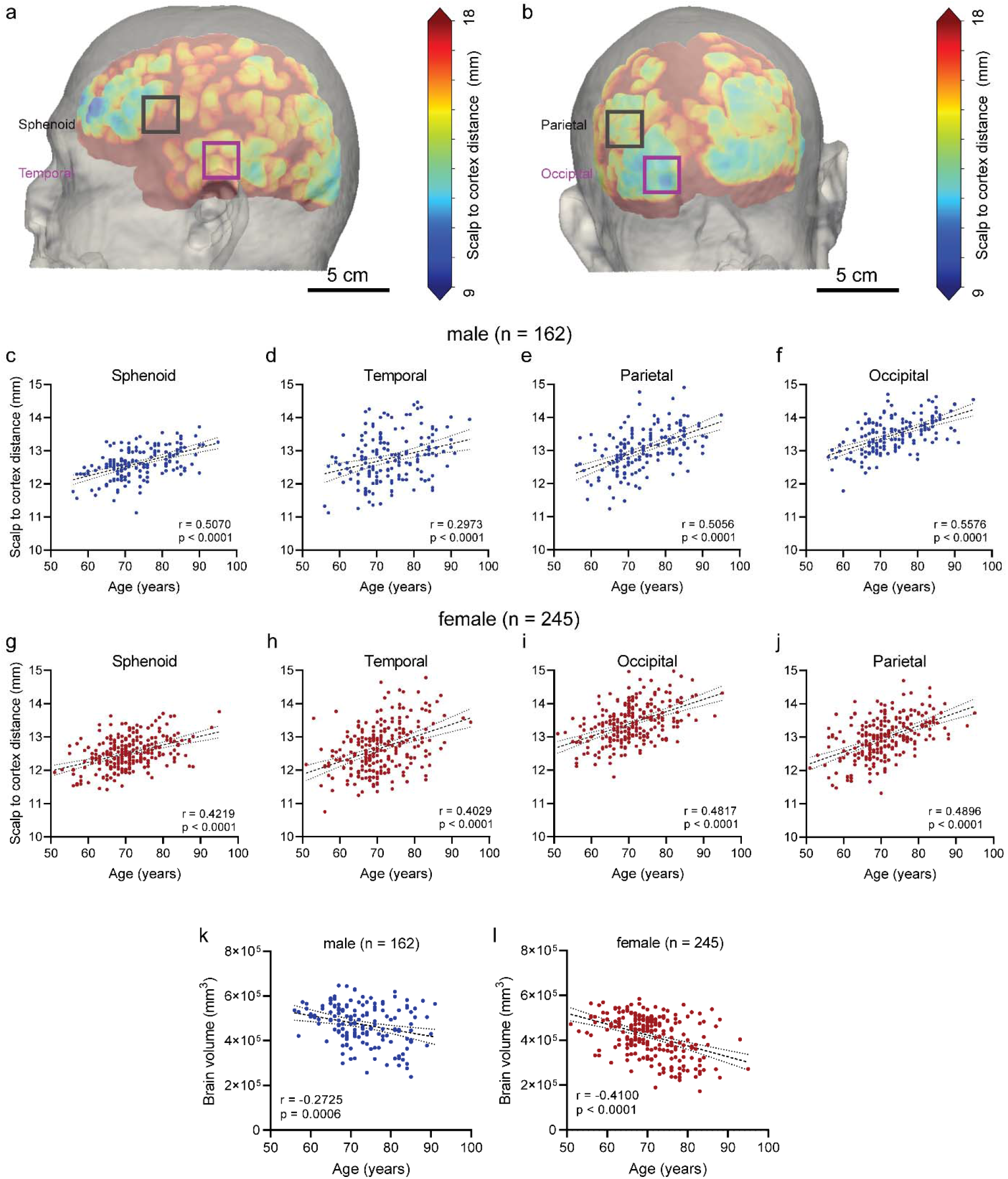
Scalp to cortex distance (SCD) map and association with age in male and female older adults. (**a**) Representative SCD map-based T_1_w MR from one 55-year-old male; gray square indicates the sphenoid cortex (a) and parietal cortex (b); magenta square indicates the temporal cortex (a) and occipital cortex (b). Scale bar = 5 cm; Color scale = 9-18 mm (blue□red); (**b-e**) Association between age and skull thickness in male participants (n = 162); (**f-i**) Association between age and skull thickness in female participants (n = 245). (**j-k**) Association of brain volume with age in male and female older adults. (j) In male participants (n = 162); (k) in female participants (n = 245).

**Table 3.** Comparison of skull thickness in male and female cognitively normal older adults of different ages.

| **Skull Thickness** | **All ages (yrs)** | **%/y** | **51-60 (yrs)** | **61-70 (yrs)** | **71-80 (yrs)** | **p** | **81-90 (yrs)** |
| --- | --- | --- | --- | --- | --- | --- | --- |
| **Male** | (n = 162) |  | (n = 8) | (n = 57) | (n = 55) |  | (n = 40) |
| SC | 1.73±0.23<br>[1.70-1.77] | 0.112% | 1.66±0.17 | 1.76±0.26 | 1.79±0.31 |  | 1.72±0.22 |
| TC | 1.65±0.21<br>[1.61-1.68] | -0.097% | 1.72±0.24 | 1.59±0.19 | 1.69±0.21 | 0.0402<sup>b</sup> | 1.67±0.21 |
| PC | 2.01±0.35<br>[1.96-2.07] | 0.146% | 1.96±0.36 | 1.96±0.37 | 2.10±0.40 |  | 2.05±0.38 |
| OC | 1.86±0.28<br>[1.81-1.90] | -0.123% | 1.97±0.59 | 1.93±0.51 | 1.95±0.51 |  | 1.90±0.33 |
| **Female** | (n = 245) |  | (n = 21) | (n = 107) | (n = 88) |  | (n = 27) |
| SC | 1.69±0.26<br>[1.66-1.72] | 0.202% | 1.53±0.23 | 1.68±0.26 | 1.75±0.27 | 0.0037<sup>b</sup> | 1.63±0.22 |
| TC | 1.61±0.20<br>[1.58-1.63] | -0.074% | 1.61±0.21 | 1.61±0.19 | 1.61±0.19 |  | 1.58±0.21 |
| PC | 1.94±0.31<br>[1.90-1.98] | 0.138% | 1.90±0.29 | 2.03±0.46 | 1.94±0.38 |  | 1.98±0.31 |
| OC | 1.75±0.18<br>[1.73-1.78] | 0.215% | 1.75±0.19 | 1.79±0.24 | 1.86±0.48 |  | 1.86±0.32 |

Data are presented as the mean ± standard deviation [confidence interval]; <sup>b</sup>, compared to the 61-70 yrs group. Two-way ANOVA with Bonferroni post hoc correction. SC, sphenoid; TC, temporal; PC, parietal; OC, occipital;

**Table 3a.**
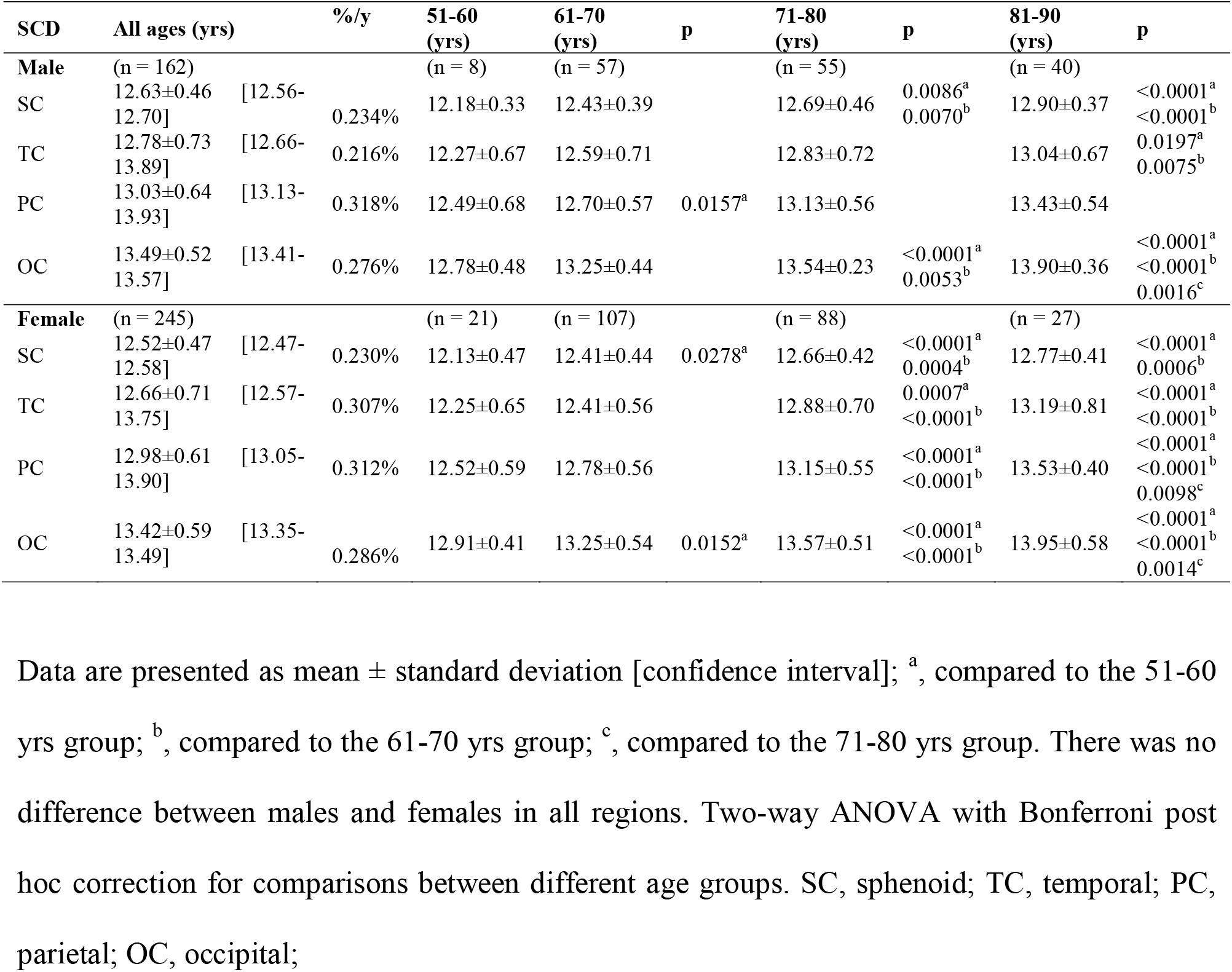
Scalp to cortex distance (SCD, mm) in male and female cognitively normal older adults of different ages.

### 3.6 Sex-dependent age-associated brain volume decrease in cognitively normal older adults

Brain volume was computed based on the cortex surface volume (**Fig. 1b**). We did not further examine the regional difference in brain volume, as there have been several well-established pipelines and cortical thickness and regional brain atrophy analysis studies previously. Group comparison of the brain volume was performed between male and female groups of all ages (Mann□Whitney) and of different age groups and between different age groups of each sex (two-way ANOVA with Bonferroni post hoc correction) (**Table 4**). Approximately 10-15% larger brain volume was observed in the male participants than in the female participants at all ages (4.58±1.09 10^5^ mm^3^, n = 162 vs 4.19±0.90 10^5^ mm^3^, n = 245; p < 0.0001), as well as in the 61-70, 71-80 and 81-90 years age groups (**Table 4**). In addition, significant brain atrophy was found in the older group compared to the relatively younger group in both female and male participants. We observed a reduced brain volume associated with age in both the male group (r = -0.3887, p < 0.0001, n = 162) and female group (r = -0.4100, p < 0.0001, n = 145) (**Figs. 4j, k**). The slope of reduction in the whole brain volume was steeper in females (-1.031%) than in males (-0.998%) (**Table 4**).

**Table 4.** Brain volume (10^5^ mm^3^) of males and females at different ages.

| Brain volume | All ages | %/y | 51-60 (yrs) | 61-70 (yrs) | 71-80 (yrs) | p | 81-90 (yrs) | p |
| --- | --- | --- | --- | --- | --- | --- | --- | --- |
| **Male** | (n = 162) |  | (n = 8) | (n = 57) | (n = 55) |  | (n = 34) |  |
|  | 4.72±0.88<br>[4.58-4.86] | -0.600% | 5.24±0.47 | 4.90±0.78 | 4.66±0.84 |  | 4.36±1.04 | 0.0059<sup>a</sup><br><0.0001<sup>b</sup><br>0.0164<sup>c</sup> |
| **Female** | (n = 245) |  | (n = 21) | (n = 107) | (n = 88) |  | (n = 27) |  |
|  | 4.19±0.90<br>[4.08-4.30] | -1.031% | 4.71±0.72 | 4.46±0.68 | 3.96±0.95 | 0.0113<sup>a</sup><br>0.0025<sup>b</sup> | 3.56±1.03 | 0.0004<sup>a</sup><br><0.0001<sup>c</sup> |
| **p** | <0.0001 |  |  | <0.0001 | <0.0001 |  | 0.0134 |  |

Data are presented as mean ± standard deviation [confidence interval]; <sup>a</sup>, compared to the 51-60 yrs group; <sup>b</sup>, compared to the 61-70 yrs group; <sup>c</sup>, compared to the 71-80 yrs group.

## 4 Discussion

Here, we developed an automatic pipeline for skull thickness and SCD computation and demonstrated the influence of age and sex on these measures in 407 cognitively normal older adults. Given the diversity in biological measures, a larger cohort is needed to evaluate the changes in skull thickness in association with sex and aging. This is the first study with a large structural MR sample size on human skull geometry in living older adults. Most previous studies are based on CT in cadaver samples and CT/MR in a small sample size of living participants. Although CT imaging is better in studying skull features, it involves radiation to living participants, which is not optimal. Moreover, structural imaging on 3 T MR provided insights into brain atrophy information in addition to skull feature information at the same time.

For T_1_w MRI head segmentation/skull stripping, various toolboxes, including FSL BET2 [30, 31], SPM12-based unified segmentation [32], BS skullfinder [20], AFNI 3dSkullStrip (3DSS) [35], ROAST [2], CHARM [19], SimNIBS 2.1 [36], ROBEX [37], BEaST [38], optiBET [39], MONSTR [40], SynthStrip (in FreeSurfer) [41], and deep learning-based approaches [42-44], have been developed. These pipelines have been very useful for brain volumetric measurements [45], automatic generation of realistic head models and electroencephalography field computations with applications in neuroimaging (e.g., diffuse optical tomography) and transcranial brain stimulation [46]. The uniqueness of our pipeline lies in the skull thickness computation and visualization in addition to the SCD outputs, which are not available in the current various packages. We showed the compatibility of our analysis with both BS- and FSL-based preprocessing. A previous study showed that FSL BET2 and SPM12 achieved better skull segmentations than BS [20]. In our study, we found great similarity between the different approaches in segmenting the skull and generating scalp mesh at the global level. However, a mismatch was observed in the automatically processed data based on FSL BET segmentation in the regions important in neurostimulation studies, such as the temporal bone and occipital bone. These results suggest that for automatic computation of skull thickness without manual correction, BS is better suited than the other two. For SCD, manual measurement has been possible [47] using the BrainRuler pipeline [48]. Our method enables efficient computation and visualization of the SCD of the whole brain in 3 minutes (whole pipeline 8 minutes).

Since we recorded the minimum regional skull thicknesses in cognitively normal older adults, the values were lower than the mean skull thickness reported in previous studies using CT [18] and MR scans in living cases [49]. Postmortem CT scans of 604 individuals showed a mean thickness of approximately 2 mm in the temporal and occipital bone [50]. Intersubject variability in the skull measures also suggests the importance of a large sample size [51]. In addition, we found that minimum skull thickness was associated with age in the sphenoid bone in females. Earlier MR studies have demonstrated the influence of skull thickening, particularly the inner skull table, associated with age and the associations between skull thickening and cognition and brain atrophy in older adults[52, 53]. A previous CT study in 120 participants aged 20-100 years indicated a nonsignificant trend of increase in the full skull thickness due to an increase in the thickness of the diploic layer [18]. Another CT skull cadaver study showed an age-related increase in skull thickness [54].

Moreover, we observed sex differences in the skull MR measures in aging cognitively normal older adults (n = 407). We found that the %skull thickening/y was higher in the sphenoid bone and occipital bone of females than in those of males. In the temporal bone and parietal bone, the rates of change were lower in the female and male groups. A previous MR study in 60 older adults (48% females) aged 71–74 years showed more inner table skull thickening in females (8.3%) than in males (6.2%) [52]. Earlier cadaver CT studies also showed intersex differences and dimorphism in the structural properties of aging femora, indicating differential bone fragility [55-57].

Here, we found that SCD increased with age in both male (n = 162) and female (n = 245) cognitively normal older adults. The increase in SCD was more rapid in the female group (0.307%/y) than in the male group (0.216%/y) in the temporal cortex, indicating faster regional brain atrophy. The rates of increase in SCD were comparable in other regions in the female and male groups (**Table 3, Fig. 4**). The temporal cortex is important in cognitive function, e.g., conceptual categorization and semantic and language processing, and is impaired early in neurodegenerative diseases such as AD [58]. Earlier lifespan studies showed brain atrophy and thinning of the cerebral cortex in aging [1, 59, 60]. An earlier study showed that the average annualized rate of hippocampal volume loss among controls was -1.55%/y [61]. Another structural MR-based study found age-related increased SCD in the left dorsolateral prefrontal cortex and not in the primary motor cortex [62].

We found a negative correlation between brain volume and age in both the female and male groups. The reduction in brain volume based on the brain surface map was more pronounced in the female group (-1.031%/y) than in the male group (-0.998%/y). An earlier CT study reported a significant relationship between cortical thinning and age for both inner and outer tables of the frontal, occipital, and parietal bones ranging between a 36% - 60% decrease from ages 20 to 100 years in females, whereas males exhibited no significant changes [18]. ICV is commonly used as a marker of premorbid brain size in neuroimaging studies, as it is thought to remain fixed throughout adulthood. Inner skull table thickening that occurs with aging would affect the ICV measure and could mask actual brain atrophy [53, 63]. Sex and age differences in the volume of regional gray matter in the normal adult human brain have been reported by using in vivo structural MR [64], as well as in postmortem examination [65]. Earlier studies showed that young females have a larger volume of cortical gray matter after correction for total brain volume [66, 67] compared with males.

There are several limitations in this study. 1) With the development of amyloid-beta and tau positron emission tomography, cellular correlates of cortical thinning are being better understood [68]. Brain atrophy is detectable in Aβ [69] and tau accumulation [70, 71] in cognitively normal older adults. However, the Aβ and tau status of the controls are not known/not analysed. 2) Since T1w MR data are being utilized for analysis, whether there are changes in body density or thickening of the inner skull table or outer table is not attainable. Further analysis using data from MR bone imaging with the zero TE sequence will provide more comprehensive information on the skull features [72, 73]. 3) Here, we used only T1w MR data for skull thickness computation. Multicontrast MRI data, T_1_+T_2,_ may provide higher accuracy; however, T_2_ MR data are not available at ADNI for the majority of cases and are thus not included in this study. 4) We did not perform ICV correction for the brain volume measurement.

## Conclusion

We developed an open-source skull thickness and SCD analysis pipeline for structural MR brain scans and demonstrated the association between skull thickness and SCD with age. The automatic efficient computation toolbox for skull thickness and SCD map analysis is potentially useful for personalized neurostimulation planning.

## Author Contributions

RN designed the study; JZ wrote the code; JZ and RN performed the analysis; VT, AG, JZ, and RN interpreted the data; and JZ and RN wrote the draft. All authors read and approved the final manuscript.

## Acknowledgements

Data collection and sharing for this project was funded by the ADNI (National Institutes of Health Grant U01 AG024904) and DOD ADNI (Department of Defense award number W81XWH-12-2-0012). ADNI is funded by the National Institute on Aging, the National Institute of Biomedical Imaging and Bioengineering, and through generous contributions from the following: AbbVie, Alzheimer’s Association; Alzheimer’s Drug Discovery Foundation; Araclon Biotech; BioClinica, Inc. ; Biogen; Bristol-Myers Squibb Company; CereSpir, Inc. ; Cogstate; Eisai Inc. ; Elan Pharmaceuticals, Inc. ; Eli Lilly and Company; EuroImmun; F. Hoffmann-La Roche Ltd. and its affiliated company Genentech, Inc. ; Fujirebio; GE Healthcare; IXICO Ltd. ;Janssen Alzheimer Immunotherapy Research & Development, LLC. ; Johnson & Johnson Pharmaceutical Research & Development LLC. ; Lumosity; Lundbeck; Merck & Co., Inc. ; Meso Scale Diagnostics, LLC. ; NeuroRx Research; Neurotrack Technologies; Novartis Pharmaceuticals Corporation; Pfizer Inc. ; Piramal Imaging; Servier; Takeda Pharmaceutical Company; and Transition Therapeutics. The Canadian Institutes of Health Research is providing funds to support ADNI clinical sites in Canada. Private sector contributions are facilitated by the Foundation for the National Institutes of Health (www.fnih.org). The grantee organization is the Northern California Institute for Research and Education, and the study is coordinated by the Alzheimer’s Therapeutic Research Institute at the University of Southern California. ADNI data are disseminated by the Laboratory for Neuro Imaging at the University of Southern California.

